# A self-exciting point process to study multi-cellular spatial signaling patterns

**DOI:** 10.1101/2020.11.04.368001

**Authors:** Archit Verma, Siddhartha G. Jena, Danielle R. Isakov, Kazuhiro Aoki, Jared E. Toettcher, Barbara E. Engelhardt

## Abstract

Multi-cellular organisms rely on spatial signaling among cells to drive their organization, development, and response to stimuli. Several models have been proposed to capture the behavior of spatial signaling in multi-cellular systems, but existing approaches fail to capture both the autonomous behavior of single cells and the interactions of a cell with its neighbors simultaneously. We propose a spatiotemporal model of dynamic cell signaling based on Hawkes processes—self-exciting point processes—that model the signaling processes within a cell and spatial couplings between cells. With this cellular point process (CPP) model, we capture both the single-cell protein bursting rate and the magnitude and duration of signaling between cells relative to spatial locations. Furthermore, our model captures tissues composed of heterogeneous cell types with different bursting rates and signaling behaviors across multiple signaling proteins. We apply our model to epithelial cell systems that exhibit a range of autonomous and spatial signaling behaviors basally and under pharmacological exposure. Our model identifies known drug-induced signaling deficits, characterizes differences in signaling across a wound front, and generalizes to multi-channel observations.

## Introduction

Complex life is largely characterized by multi-cellular structures (1). Classical multi-cellular processes such as the patterning of cells within a tissue and the precise spatial arrangement of tissues within an organ are the product of different gene expression programs organized carefully over space and time (2). These different programs emerge from both intra-cellular pathways governing gene and protein expression on a single-cell level and the inter-cellular signaling that allows cells near one another to interact. Understanding how these networks are regulated as well as the factors leading to their dysfunction are topics of active research (3).

*Intra-cellular signaling* is a term used to describe information-carrying modifications of proteins in a single cell. One example of this is the extracellular signal-regulated kinase (Erk), which is activated by phosphorylation in response to changes in the cell’s environment. Signaling proteins can then operate on downstream effectors such as transcription factors that regulate gene expression.

*Inter-cellular* signaling specifically involves signaling as a result of an input delivered by a neighboring cell. Often, this involves the release of ligands from one cell that bind to receptors on a neighboring cell and cause a change in behavior of the neighbor cell. Both intra- and inter-cellular signaling may make use of the same signaling protein. For example, Erk can be activated by the presence of growth factors in the surrounding media, or upon cleavage and binding of growth factors from an adjacent cell.

Nearly every cell in a physiological context is simultaneously processing information about its own state (intra-cellular) as well as the states of cells around it (inter-cellular). Therefore, these two modes of communication may interact in complex and unexpected ways, especially when they make use of the same signaling proteins. Decoupling their relative effects on cell state is challenging and often requires invasive perturbations such as pharmacological inhibitors that may have unforeseen consequences on cell or tissue health. Nevertheless, estimating the relative contributions of intrinsic cellular behavior and extrinsic spatial signaling is an important goal for understanding multi-cellular systems. This is particularly true with the advent of cellular imaging modalities that allow us to visualize signaling behaviors in single cells, heterogeneous multi-cellular ensembles, and even *in vivo* tissue (4).

Here, we focus on a case of one signaling pathway being used to convey information about both intra- and inter-cellular cell states. The mammalian Ras/Erk pathway has been found to display transient “pulses” consisting of pathway activation followed by rapid deactivation, in a range of epithelial cell types (5, 6). These pulses can be modulated by environmental context such as the presence of certain growth factors, as well as physical perturbations such as a wound (4). Moreover, cells have been found to “transmit” pulses of activity from cell to cell, as well as pulse autonomously, suggesting that the Ras/Erk pathway can operate in both intra- and inter-cellular signaling in these cell types (5, 6). Since epithelial tissues largely derive their functionality from inter-cellular communication and multi-cellular behavior, which regulates their differentiation and growth, it is likely that Ras/Erk pulses in epithelia are not simply an epiphenomenon but rather functionally linked to tissue-level function, for instance, to the ability of a cell to leave its stem cell niche and undergo subsequent differentiation (7).

Models have been proposed that approximate spatial signaling patterns to simulate signaling behaviors of cells. Turing models based on reaction-diffusion equations describe the behavior of biological systems spatially by approximating the tissue as a continuum (8, 9). Cellular automata models equip each member of a discrete set of agents with a primitive set of directives, and then observe how the resulting system of agents evolves across time (10). To our knowledge, inferring signaling parameters for these models from observations is rare, limiting their applicability to observational data from experimental systems.

In this paper, we introduce a statistical approach to modeling the pulsing times of each cell in a neighborhood as a function of their spatial organization. We treat collections of signaling pulses as realizations of a *point process* – a stochastic model of events over time or space (11). Self-exciting point processes, known as Hawkes processes, have been successfully used to model social media interactions (12, 13), financial time series (14, 15), neuron spike trains (16), and genomics events along DNA sequences (17, 18), as well as a range of other time-varying or space-varying processes (19, 20). These processes allow us to explicitly parameterize the rate of cell pulsing and to quantify the influence of a pulsing event at one cell on the probability of a pulsing event in neighboring cells in the future as a function of distance. Previous work has mostly focused on learning the connectivity of a network given data; we are interested in learning the strength of connections based on a given spatial network.

Our model —the cellular point process (CPP) — quantitatively estimates the base rate of pulsing with the rate of intra-cellular signaling jointly with the strength of inter-cellular signaling using experimental data. The CPP is an adaptation of the original Hawkes process that defines the signaling strength as a function of the distance between cells. The CPP model parameters are estimated by maximum likelihood methods using data that capture pulse times and spatial coordinates for each cell annotated in an imaging experiment. These types of experiments are becoming increasingly tractable in many laboratory environments (6). We demonstrate a correlation between the duration (i.e., number of signaling events) and scale (i.e., number of cells) quantified in an experiment, and the accuracy of the intra- and inter-cellular signaling rates inferred by the CPP. This suggests that our model can be applied to a relatively wide range of imaging data to deconvolve pulsing rates due to intra- and inter-cellular signaling patterns.

We validate the CPP’s ability to estimate the relative contributions of intra- and inter-cellular signaling on bursting rates in simulated data where these contributions are known. We then analyze mouse epidermal stem cells, or keratinocytes, which display naturally occurring Ras/Erk dynamics *in* and *ex vivo* (4, 6). We quantify the decrease in spatial signaling when these cells are treated with a known cell signaling inhibitor compared to untreated cells. Then, we examine and disentangle the inter- and intra-cellular contributions of a variety of pharmacological kinase inhibitors on Erk bursting dynamics in keratinocytes. Next, we study the contributions of inter- and intra-cellular signaling on the response of Madin-Darby Canine Kidney (MDCK) cells to an acute wounding event, finding that both factors change as a function of distance away from the wound. Finally, we demonstrate that the model estimates how multiple reporter channels interact with each other across cells.

## Model and Methods

### The Cellular Point Process (CPP) Model

Point processes are probabilistic models of events in some mathematical space, generally used to model event occurrences across space or time (or both) (19–21). A point process can be defined by the conditional intensity function, *λ*(·), which represents the expected infinitesimal rate at which events occur (22). We first describe a one-dimensional point process that starts at *t* = 0. This point process produces *N* events over an interval of time *δt* > 0. At any moment *t* > 0, there is a history of previous events, *H_t_*, that consists of the times, {*τ*_1_, *τ*_2_, …, *τ*_*i*_ < *t*}, of each previous event. The conditional intensity function is then defined as:

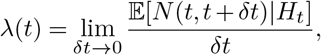

where *δt* represents a non-negative interval of time.

A simple point process (23, 24), such as a Poisson process, may have a constant conditional intensity over time:

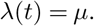

In this simple Poisson process, the expected number of events is *μ*Δ*t* over time Δ*t*. However, biological process are generally *non-stationary* — the expected number of events changes as a function of time. This non-stationary behavior comes from *self-excitation*, meaning an event makes future events more likely for a period of time. This phenomenon can be caused by a variety of mechanisms but is broadly referred to as positive feedback (25, 26). The conditional intensity function can be altered to represent such non-stationary behavior.

Hawkes processes (22, 27) model self-excitation with a conditional intensity dependent on history:

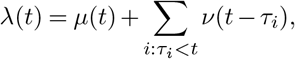

where *μ*(*t*) represents an autonomous underlying rate of events, *τ_i_* is the time of event *i* ∈ *H*_*t*_, and *ν* is a kernel function defining the association between previous events and the conditional intensity.

In the multi-variate case (13, 15, 28) where more than one set of events is observed simultaneously, the conditional intensity of one dimension *λ_j_* is a function of the history in *K* dimensions:

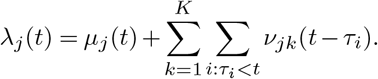

In this model, events in one variable may influence the conditional intensity function of other variables through the kernel function *v*_*jk*_(·).

To model cells from a homogeneous population recorded on a two dimensional plane, we treat each cell as a different variable in a multi-variate point process. We assume that the autonomous component *μ*_*j*_(*t*) is a positive constant *μ* > 0 over time and across cells. Based on observations from prior work (6), the time kernel *ν_jk_* is assumed to have a log-normal shape with zero mean and variance *b*_*jk*_:

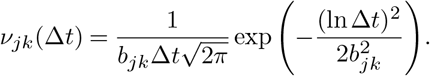

We assume a constant *b*_*jk*_ = *b* for all *j* ≠ *k* that represents inter-cell signaling and *b*_*jk*_ = *b_self_* for *j* ≠ *k* that represents intra-cell excitation. Intuitively, when a cell pulses, the conditional intensity of pulsing in its neighbors increases until peaking at time exp(−*b*^2^), after which the conditional intensity decreases back towards the baseline *μ*. Biologically, this captures the expected delay between subsequent signaling events.

We make *a_jk_* a function of the distance between cells to capture the spatial nature of cell-cell interactions:

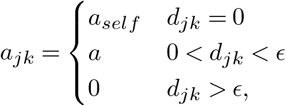

where *d_jk_* represents the distance between cells, *ϵ* is a radius inside which signaling is possible, *a* is a positive constant that quantifies the strength of cell-cell interactions, and *a_self_* is a positive constant for the magnitude of intra-cell self-excitation.

For any individual cell *j* among *K* cells, the full cellular point process (CPP) model is defined as:

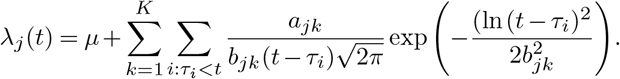

We can generalize the CPP model to account for multiple types of observations per cell, such as different fluorescence channels corresponding to different components of a protein signaling network.

In this case, each channel and protein-protein interaction has a different set of model parameters. Given *L* channels capturing different proteins, and letting *ℓ* represent a particular channel, there is:

- *μ*_*ℓ*_ > 0 for each channel, representing *L* parameters;
- *a*_*ℓ*,*ℓ*′_ > 0 for each channel, representing *L*^2^ parameters;
- *a*_*self, ℓ*_ > 0 for each channel, representing *L* parameters;
- *b*_*ℓ*,*ℓ*′_ > 0 for each channel, representing *L*^2^ parameters;
- *b*_*self, ℓ*_ > 0 for each channel, representing *L* parameters;

Thus, for any channel *ℓ*_*j*_ for cell *j*, the conditional intensity according to the CPP is:

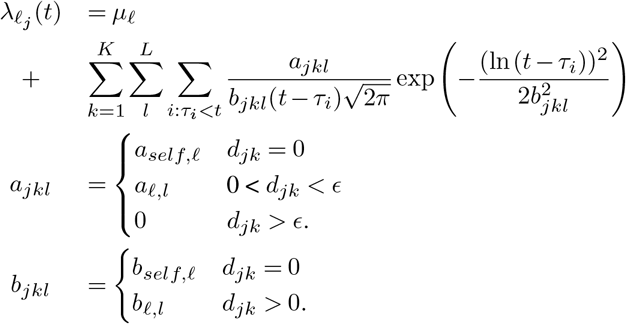

### Relationship to previous models

A number of related models have been used to address biological patterning in the past. Here, we will focus chiefly on the 2D Ising lattice model and the Kuramoto oscillator system, as these are the closest in context to the model that we propose.

The 2D Ising model is used to describe a 2-dimensional array of spins, each of which can be in one of two discrete states (spin up or spin down). This model has been implemented in creating discrete patterns in the study of reaction-diffusion systems with coupled agents (29). The operator function that gives the energy of the system as a function of the spin states *σ* of all of the constituent particles is called the *Hamiltonian*:

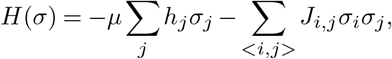

where the first term *μ* Σ_*j*_ *h*_*j*_*σ*_*j*_ is the effect of an external magnetic field *h* on a particle’s spin state, weighted by *μ*, and the second term 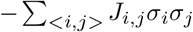 is the coupling term *J* between spins.

In our model, we can think of the coupling terms as similar to those presented in our model; i.e., cells interact with other cells within a small neighborhood around them. The magnetic field term is usually held as constant over the system, and is similar to our term *μ* that represents the probability of randomly pulsing in a cell-autonomous fashion. The Hamiltonian for the 2D Ising model is therefore similar in spirit to our model as described.

However, as many studies have shown positive feedback effects that give rise to self-excitation processes in cells, this model does not capture a reasonable range of cell signaling behavior. Additionally, upon examining sheets of cells experiencing Ras/Erk pulses, we rarely see “stable states” of cells that are constantly on or off, in contrast to the steady state behaviors often seen in Ising model simulations.

A second model used for coupled cells, and in particular cells capable of oscillations, is the Kuramoto model (30). The Kuramoto model treats cells as entities having an intrinsic oscillator “frequency” from a continuous range of values. Cells are also coupled to some number *N* of adjacent cells, with a “coupling constant” *K*:

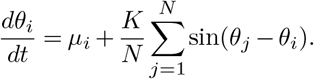

Here, the intrinsic frequency contribution *μ_i_* describes the cell-intrinsic change in oscillator phase *θ_i_*. This is akin to our *μ* parameter. In this function, the contribution of adjacent cell states to the state of cell *i* is distinct from the contribution of the autonomous behavior of cell *i*. Since the state of each cell is in the space of oscillator frequencies, this parameter is forced to oscillate through the sin of the difference in frequencies rather than parameterizing a random process, making the effects of neighboring cells periodic and therefore deterministic instead of stochastic. This deterministic function precludes transient signaling events from occurring. The Kuramoto model, when simulated for long time courses, has been shown to relax into smooth patterns of phases with clear boundaries between regions of opposite phase (30); however, this is not the signaling pattern that we are trying to capture in pulsatile cells.

While our CPP model incorporates some of the elements of these two related models, it is more suited to capture the signaling patterns we observe in experimental data. In the design of the CPP model, we make use of two terms that describe cell-autonomous and cell-to-cell signaling contributions. In all three cases, cell-to-cell signaling terms are applied over a small region around the cell in question; for example, in the 8-cell neighborhood of a square in a 2D lattice. The range of states that a cell can occupy differs among the three models. While the Ising model and our CPP model capture binary states (on or off), cells in the Kuramoto oscillator model may take values over a range of oscillator frequencies. These values, however, have the downside of being modeled deterministically, without allowing for the possibility of stochastic events.

The CPP model makes use of discrete states– more specifically, we model the bursting state of a cell as being on or off with respect to a specific protein. This point process approach does not allow for different amplitudes of signaling, but does take advantage of a framework that allows for phenomena such as positive feedback that create self-excitatory pulse trains of signaling pathway activation. Moreover, the structure of the signaling term in the CPP model allows us to take into account the history of a cell’s signaling state, which is not a part of the Ising or Kuramoto models, and allows for a more explicit treatment of self-excitatory processes in biological systems.

Both the Ising and Kuramoto models approach a “steady state” pattern of phases or spins as time goes to infinity. However, active behavior and constant emergence of non-stationary fluctuations in a population limit the applicability of these models to data. A stochastic, self-exciting, and history-dependent model such as the CPP described here better captures these behaviors.

### Inference for the CPP

The conditional intensity *λ_j_*(*t*) of the CPP has six parameters to be inferred: *a*, *a_self_*, *b*, *b_self_*, *μ*, and *ϵ*. Biologically, Erk signaling is regulated across cells by interactions between membrane proteins, limiting the signal to neighboring cells. Since the (*x, y*) spatial coordinates of each cell in our experimental data represent the cell center, we set *ϵ* to 60 pixels, corresponding to roughly 84 *μm*, unless otherwise stated. This value was obtained through visual inspection of the raw imaging data. We noticed that cells on average had 5 neighbors in direct contact with them, forming a local neighborhood of cells around each cell. We then chose *ϵ* such that the average cell had 5 neighbors. Of course, this parameter is calibrated to the cell distributions observed in our experiments; in future applications it will need to be adapted for images with different resolutions or cell geometries.

We optimize the remaining parameters by maximizing the log likelihood of the observations. Given a history *H*_*t*_ with *N* events {*τ*_1_, *τ*_2_, …, *τ*_*n*_} in time interval [0, *T*], the log likelihood is (12):

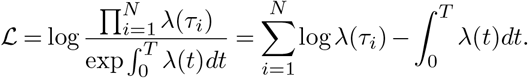

We maximize this likelihood using automatic differentiation from PyTorch (31). We use stochastic gradient descent (SGD) to minimize the negative log likelihood, with stopping criteria when a local minimum has been reached (the negative log likelihood at iteration *i* is greater than the negative log likelihood at iteration *i* − 1) or when the absolute change in log likelihood between iterations is less than 0.001%.

### Simulations

We simulate histories of peaks in one channel from the generative model to test whether inference accurately estimates the true parameters. We first generate 50 sets of {*μ*, *a*, *a*_*self*_, *b*, *b*_*self*_}. Each parameter is selected from a uniform distribution with set minimums and maximums (Table 1). The maximum distance of spatial interactions, *ϵ*, is kept constant at 60 pixels, corresponding to roughly 84 *μm*. For each parameter set, for every combination for cells in [50, 75, 100, 125, 150, 175, 200] and number of peaks in [100, 250, 500, 1000, 2500], a history is simulated. Cell positions are drawn from a uniform distribution between (0, 0) and (*x_max_, y_max_*). To match data collected from microscopy, *x_max_* = *y_max_* to create a square region. The maximum coordinates as well as *ϵ* are set such that the expected number of neighbors, 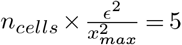, similar to 2data from Goglia et al. (6). This corresponds to an inter-cellular distance of approximately 30-50 *μm*. For each history, we fit the model until convergence. We evaluate the goodness-of-fit by calculating the mean squared error (MSE) between estimated and true parameter values.

**Table 1.**
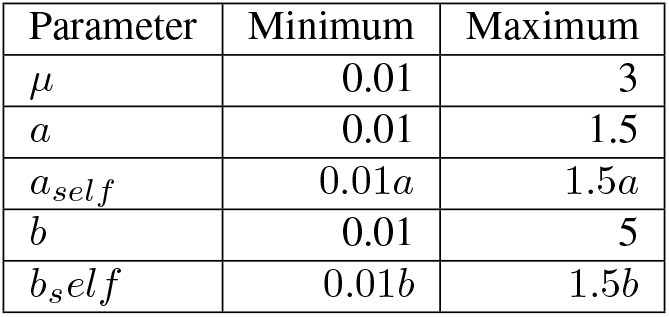
Parameter range for simulating a spatial point process.

| Parameter | Minimum | Maximum |
| --- | --- | --- |
| $\mu$ | 0.01 | 3 |
| $a$ | 0.01 | 1.5 |
| $a_{self}$ | $0.01a$ | $1.5a$ |
| $b$ | 0.01 | 5 |
| $b_{self}$ | $0.01b$ | $1.5b$ |

## Experimental Methods

### Cell culture and generation of transgenic cell lines

Dorsal epidermal keratinocytes derived from CD1 mice and stably expressing a lentivirally-delivered histone H2B-RFP and ErkKTR-BFP (6) were cultured as described previously (32). Briefly, keratinocytes were grown in complete low calcium (50 mM) growth media (‘E media’ supplemented with 15% serum and 0.05 mM *Ca*^2+^) in Nunclon flasks with filter caps (Thermo-Fisher) and were maintained in a humidified incubator at 37 C with 5% *CO*_2_. Cell passage number was kept below 30. Keratinocyte media was prepared as per prior work (32).

To create pFos-GFP expressing cells, dorsal epidermal keratinocytes were derived from TARGATT mice containing a safe harbor locus with an attB insertion site (Applied Stem Cell, Milpitas, CA). A vector containing the minimal Fos promoter driving a destabilized GFP and a CMV promoter driving a hygromycin resistance gene was constructed using Infusion cloning, and flanked with attP sites for insertion into the attB sites in TARGATT keratinocytes. Keratinocytes were cotransfected with this reporter plasmid as well as a plasmid encoding the phiC integrase driven by a CMV promoter, which, when expressed, completed the integration of the reporter construct into the safe harbor locus.

Cells were selected for expression with hygromycin (Sigma Aldrich). Prior to imaging experiments, cells were transduced with lentiviral vectors encoding a H2B-RFP marker, as well as with ErkKTR-iRFP.

Imaging experiments were performed in 96-well black-walled, 0.17mm high performance glass-bottom plates (Cellvis). For plating cells, wells were pre-treated with a solution of 10 mg/mL bovine plasma fibronectin (Thermo Fisher) solubilized in phosphate buffered saline (PBS) to support cell adherence. Two days before imaging, keratinocytes were seeded at approximately 96, 000 cells/well in 100 uL of low-calcium E media (in a 96-well plate). Glass-bottomed plates were briefly centrifuged at 800 rpm to ensure even plating distribution, and cells were allowed to adhere overnight. 24 h before imaging, wells were washed 2-3x with PBS to remove nonadherent cells and were shifted to high-calcium (1.5 mM *CaCl*_2_) complete "E" media to promote epithelial monolayer formation. For experiments in growth factor-free (starvation) media, cells were washed once with PBS and shifted to high-calcium P media (DMEM containing only pH buffer, penicillin/streptomycin, and 1.5 mM *CaCl*_2_) eight hours before imaging. To prevent evaporation during time-lapse imaging, a 50 mL layer of mineral oil was added to the top of each well immediately before imaging.

Imaging was performed on a Nikon Eclipse Ti confocal microscope, with a Yokogawa CSU-X1 spinning disk, a Prior Proscan III motorized stage, an Agilent MLC 400B laser launch containing 405, 488, 561, and 650 nm lasers, a cooled iXon DU897 EMCCD camera, and fitted with an environmental chamber to ensure cells were kept at 37 C and 5% *CO*_2_ during imaging. All images were captured with a 20X air objective and were collected at intervals of 3 min.

For TAPI-1 experiments, drug was obtained from SelleckChem (Houston, TX) and diluted to 10x the relevant concentrations in DMSO. 11 *μL* of drug was added to 100 *μL* of cells in 96 well plates immediately before imaging. For drug treatment experiments (Fig. 4), drugs were added to a final concentration of 2.5 *μM*.

For MDCK wound healing experiments, MDCK cells were maintained in minimal essential medium (MEM; ThermoFisher Scientific, 10370-021) supplemented with 10% fetal bovine serum (FBS; Sigma, 172012-500ML), 1x Glutamax (ThermoFisher, 35050-061), and 1 *mM* sodium pyruvate (ThermoFisher,11360070), in a 5% *CO*_2_ humidified incubator at 37C. For time-lapse imaging, MDCK cells were plated on 35 mm glass-base dishes (Asahi Techno Glass). Before time-lapse imaging, the medium was replaced with FluoroBrite (Invitrogen) supplemented with 5% FBS and 1x Glutamax.

For the generation of MDCK cell lines stably expressing the FRET biosensor, a PiggyBac transposon system was used (5, 33). The pPBbsr-based FRET biosensor and pCMV-mPBase (neo-) encoding the piggyBac transposase were co-transfected into MDCK cells using an Amaxa nucleofector system (Lonza, Basel, Switzerland) at a ratio of 4:1. The cells were selected with 10 mg/ml of blasticidin S for at least 10 days. Singe cell clones expressing EKAREV-NLS were further isolated by limited dilution.

MDCK cells (4 × 10^5^ cells) were plated on 35 mm glass-based dishes. Two days after seeding, confluent cells were scratched with a 200 *μL* pipette tip to establish the wound. Just after scratching, the media and dislodged cells were aspirated and replaced by FluoroBrite with 5% FBS and 1x Glutamax. Immediately after replacing the media, the cells were imaged with an epi-fluorescence wide-field microscope. The cells were imaged every three minutes for 12 hours. DMSO, 100 nM trametinib, or 10nM TAPI-1 was added two hours after starting time-lapse imaging.

### TAPI Dose Response

A number of publications on the phenomenon of spatially coupled Ras/Erk pulses have noted that the matrix metalloprotease inhibitor TAPI-1 is capable of reducing the extent of cell-to-cell signaling in pulsatile activity (5, 34). The drug inhibits the cleavage and release of ligands that activate the Ras/Erk pathway in adjacent cells, a process called juxtacrine signaling(35). This body of work suggests that TAPI-1 specifically inhibits inter-cellular, but not intra-cellular, signaling pulses.

Previous work on TAPI-1 as an inhibitor of spatial signaling in pulsatile activity has been limited to analyses of approximately 5 10 cells at a time, and focuses on isolated instances of cells losing spatial coupling upon TAPI-1 addition, rather than on a population level response. We treated keratinocytes with a range of TAPI-1 doses, at 5, 10, and 20 *μ*M, and imaged cells from the point of TAPI-1 exposure. An untreated well to which only the solvent dimethylsulfoxide (DMSO) had been added was imaged as a vehicle control, since TAPI-1 was solubilized in DMSO prior to addition to the well. DMSO has not been found to affect Ras/Erk activity dynamics (6). Imaged cells were incubated in growth factor-free media, and cells were imaged every three minutes for 12 hours after the addition of TAPI-1. We noticed that cells went through a period of deactivation after the addition of the drug after which pulsing resumed; to remove this from our analysis, time-series were truncated to the last 6 hours of imaging. Time-series measurements were converted to a series of peaks for each cell as described previously (6). The model was fit for each well separately until convergence.

### Estimating the effects of different drugs on keratinocyte signaling

We next fit the model to data from prior work (6), in which keratinocytes were treated with various receptor tyrosine kinase inhibitors (RTKi), which target proteins upstream of endogenous Erk activity. The data consist of 450 wells with 432 different drug treatments and 18 DMSO vehicle controls (which contain no inhibitor) (see Supplemental Material). Imaged cells were incubated in growth factor-free media, and RTKi was added 30 minutes prior to imaging. Cells were imaged every three minutes for 12 hours after addition of RTKi. Time-series measurements were converted into a series of peaks for each cell as described previously (6). The model was fit to data from each well separately until convergence. Due to differences in spatial organization across wells, the signaling radius *ϵ* was set independently for each well such that each cell had on average five neighbor.

### MDCK wound healing

Extensive prior work has been done on the association between Ras/Erk pathway activity and cell proliferation and migration, events that are critical for regeneration and wound healing. In light of this, we demonstrated the use of live-cell Ras/Erk activity reporters in combination with our model to characterize the behavior of the Ras/Erk pathway in response to an acute wounding event. Since a wound has a particular spatial location relative to different cells, we used our model to quantify signaling rates at various distances from the wound. To do this, we collected data on a large sheet of Madin-Darby Canine Kidney (MDCK) cells, which are widely used for studies of collective cell motility. Cells expressing the EKAREV Erk activity reporter(36) were established using a piggyBac transposon system (33, 37) and sorted to ensure uniform expression of the reporter construct. For wound healing assay experiments, a wound was inflicted on cells by scratching a pipette across a confluent layer of cells, and the sheet of cells was imaged every three minutes for 12 hours. Nuclei were segmented, and Erk activity was measured for each cell over time using the cell tracking software TrackMate (6). Due to cell movement over the course of the experiment, the field of cells was split into ten bins according to each cell’s distance from the wound edge along the x-axis at the start of the experiment, immediately after the wound had been inflicted. Our model was fit until convergence to each individual bin, consisting of the cells present in that spatial bin at the first time point, over the duration of the wound healing process. As a control, we also binned cells along the y-axis, to ensure that these ten bins result in identical estimated signaling behaviors since these bins run parallel to the wound. Data collected in the presence of the matrix metalloprotease inhibitor TAPI-1 and the MEK inhibitor Trametinib were also processed and analyzed in the same manner.

### Analyzing behavior in multiple channels

As described earlier, the CPP model makes it possible to examine couplings between separate channels, for example, to analyze separate components of a signaling network. The Ras/Erk pathway has a well-defined set of target genes, called Immediate Early Genes (IEGs), that respond acutely and rapidly to Ras/Erk stimulation. We engineered mouse keratinocytes to express a destabilized green fluorescent protein (dGFP) with a half-life of ~ 1 hour, under the control of the minimal promoter of the IEG cFos. Using these cells, we could measure Erk-KTR as well as dGFP across 24 hours in the same cells. Time-series measurements were converted into a series of peaks for each channel in each cell (6). CPP was fit run until convergence for each experiment to estimate the *μ, a, a_self_, b,* and *b_self_* terms for each channel and cross-channel interaction.

## Results

The cell point process (CPP) model treats each peak in a cell as an event in a self-exciting point process where events in one cell influence the likelihood of an event occurring in the future in that cell and in neighboring cells (Figure 1). The CPP model estimates five parameters from a list of events in different cells with known spatial organization:

- *μ*, the baseline, autonomous frequency of events;
- *a*, the strength of signal each event emits to the cell neighborhood. Higher *a* indicates stronger signaling effects;
- *a_self_*, the strength of signal each events emits to self-excite the cell of the event. Higher *a_self_* indicates stronger intra-cellular effects;
- *b*, the variance of a log normal kernel that defines the effect of any event on the conditional intensity of its neighbors. Higher *b* corresponds to larger variance in the time between events; and
- *b_self_*, the variance of a log normal kernel that defines the effect of any event on the conditional intensity of its own cell.

**Fig. 1.**
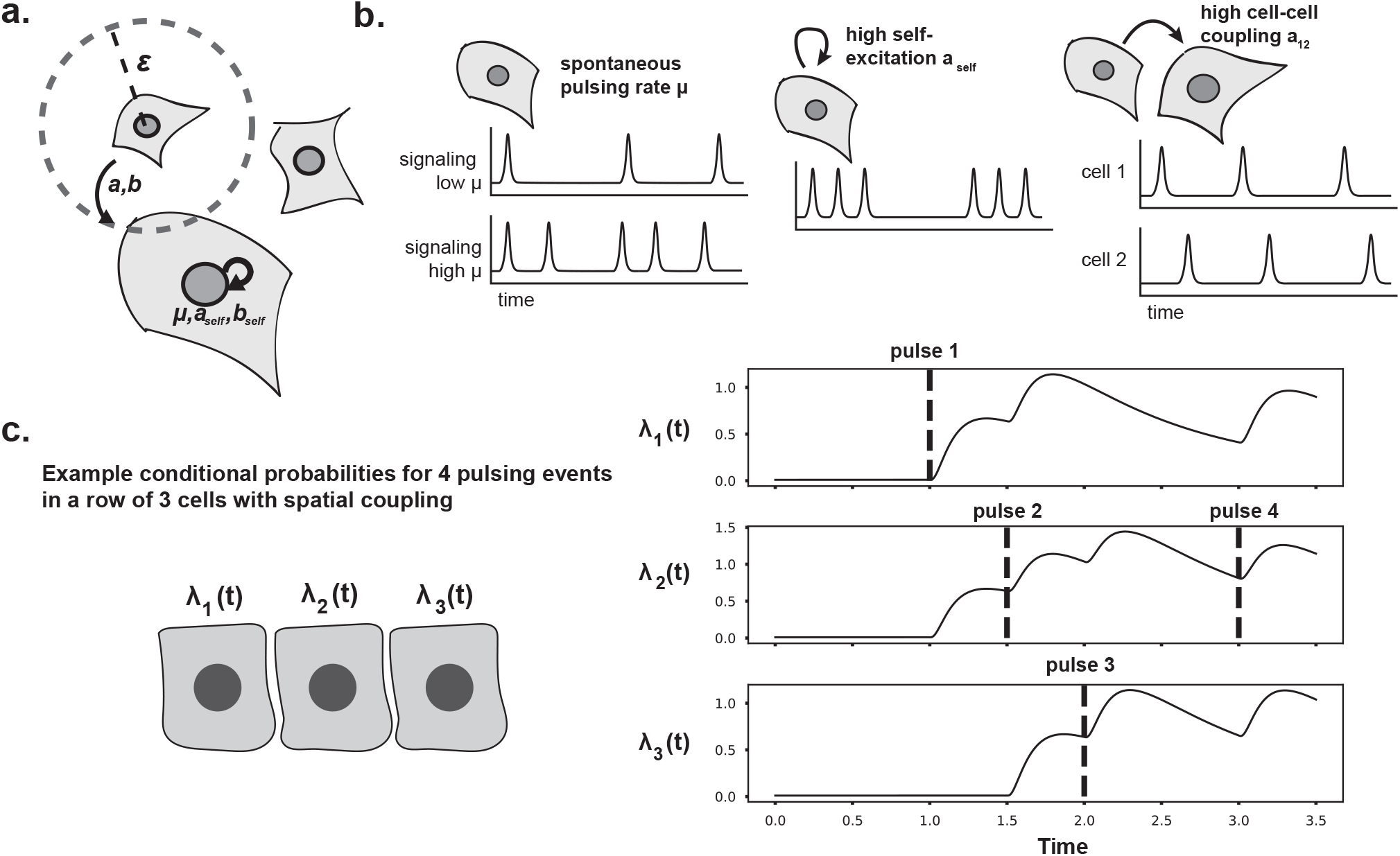
(a) Three cells demonstrating autonomous pulsing *μ*, self signaling effects parameterized by *a_self_, b_self_*, and cell-proximity specific effects within some radius *ϵ*, with magnitude of those effects parameterized by *a, b*. (b) Effect of the parameters in (a) on signaling pulses in cells. (c) Sample conditional intensity (*λ*(*t*)) plots for 3 cells, where cell 1’s initial pulse increases the chance of pulsing for cell 3 and itself through spatial coupling. High intensity means more expected pulses.

We first validate the CPP model’s ability to identify spatial and autonomous signaling on simulated data where these factors are known *a priori*. We then turn to a naturally occurring and experimentally tractable system of mouse epidermal stem cells, or keratinocytes, that display naturally occurring Ras/Erk dynamics *in* and *ex vivo* (4, 6). We quantify the natural inter-cellular and intra-cellular signaling effects in these cells, and validate the CPP model by quantifying decreases in spatial signaling when cells are treated with a known signaling inhibitor. Next, we quantify the spatial and autonomous signaling effects of a variety of drugs on keratinocytes from a prior assay (6). We also quantify the spatial and autonomous signaling effects of three drugs on cell behavior during wound healing stratified by distance from the wound (34). Finally, we demonstrate that the model can estimate parameters from more complex histories with multiple channels per cell using data with two fluorescence channels from mouse keratinocytes.

### Spatial point processes estimate parameters from simulated data

We first verify that the model accurately estimates parameters from simulated data with known ground truth. Observations were simulated from the generative model across a range of observed cells and total number of observed events. We find that the CPP model is able to accurately estimate the parameters of the generative model with low MSE (Table 2; Figure 2). A control estimate of *μ*, the average number of peaks per unit time per cell, has a higher error (Average MSE = 1.22) than the CPP model’s estimates.

**Table 2.** Mean squared error (MSE) of CPP-estimated parameters to true ground truth parameters from simulated data.

| Parameter | MSE |
| --- | --- |
| $\mu$ | 0.037 |
| $a$ | 0.12 |
| $a_{self}$ | 0.068 |
| $b$ | 4.11 |
| $b_{self}$ | 1.38 |

**Fig. 2.**
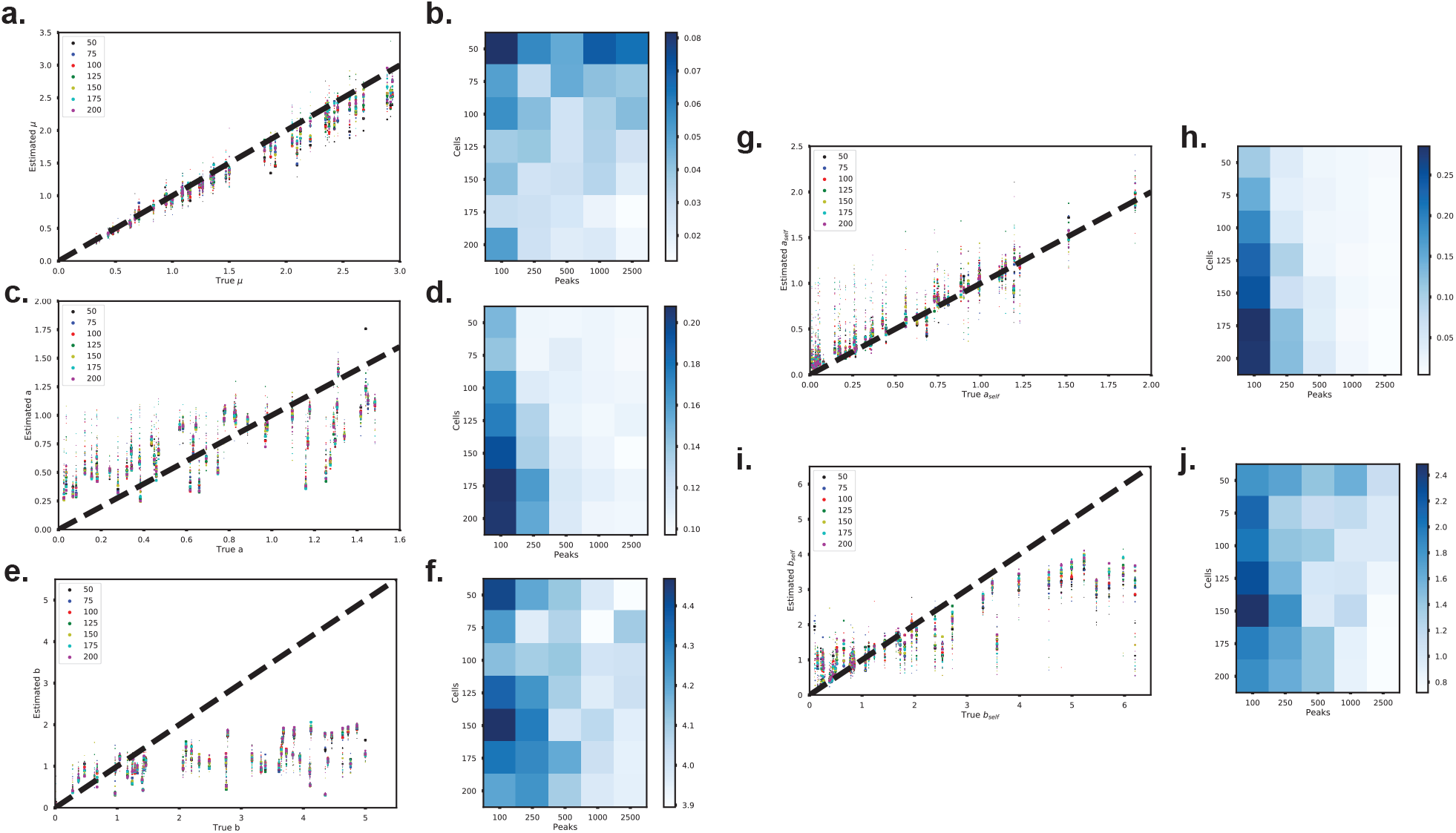
(Left) Scatter plots of true versus estimated (a) *μ* (c) *a* (e) *b* (g) *a_self_*, and (i) *b_self_*. Dot color represents the number of cells and dot size represents the number of events for that simulation. (Right) Heatmap of mean squared error (MSE) of estimated (b) *μ* (d) *a* (f) *b* (h) *a_self_* and (j) *b_self_* for different simulation parameters number of cells and number of peaks.

The accuracy of the parameter estimates varies over the number of cells and number of events observed. We observed that estimates for *μ* are worse when the number of cells is low (Figure 2b). The estimates for signaling parameters *a*, *a_self_*, *b* and *b_self_*, in contrast, degrade as the number of observations decreases (Figure 2d,f,h,j). We found that *a* is usually overestimated when there are small numbers of peaks (Figure 2c). The accuracy of estimates also depends on the true parameter values. The autonomous parameter *μ*, and intra-cellular signaling duration parameter *b_self_* tend to be underestimated when their true value is larger (Figure 2a,e,i). Estimation for inter-cellular signaling duration parameter *b* is less accurate than the other parameters (Table 1), so we ascribe less significance to it’s estimate in later sections. Nevertheless, the small errors of the parameter estimates demonstrates that the CPP model accurately estimates signaling parameters from experimental data. The data we collected has around 150 cells and 1000 peaks, a region where CPP’s estimates for each parameter are close to the true value from simulations.

### Spatial point processes capture the effect of pharmacological treatments on keratinocyte behavior

Next, we evaluate the CPP model’s ability to identify changes in signaling behavior across pharmacological treatments of cells. Mouse basal keratinocytes are epidermal stem cells that form monolayers *in* and *ex vivo*. Studies have shown dynamic signaling behavior in the Ras/Erk pathway of these cells linked to spatial patterning (5). TNF-*α* protease inhibitor (TAPI-1) is a matrix metalloproteinase inhibitor (38) that prevents spatial signaling and Erk activation between cells (5). We used a KTR fluorescent marker (39) to image Erk concentration over time across sheets of keratinocyte cells dosed with 5, 10, and 20 *μ*M of TAPI-1, as well as a control group of untreated cells.

Our CPP model is able to quantitatively capture the change in keratinocyte signaling behavior as a function of TAPI-1 concentration (Figure 3). We find that the estimated autonomous 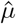 parameter and strength of spatial signaling 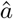 decrease with increased TAPI-1 concentration (Table 3). We also observe that the self-exciting parameter *a_self_* does decrease but only about 20%. We would not expect TAPI-1 to change the self-excitation signaling of keratinocytes. The kernel parameter 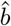 and 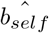 increase with increased TAPI-1 concentration, representing an increase in the time between peaks (Table 3). We note that the values for 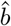 and 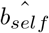 are the same; this is due to the inference methods. We estimate *b_self_* as a multiplier of *b*, initialized at one. The nearly identical values indicate that the gradient updates for the *b_self_* multiplier were small given these data. This finding highlights an important advantage of our model. In contrast to simpler models such as the Ising model, it allows for calculating an explicit term for memory—i.e., the propensity for a cell to change state as a function of state changes that have occurred in its recent history.

**Fig. 3.**
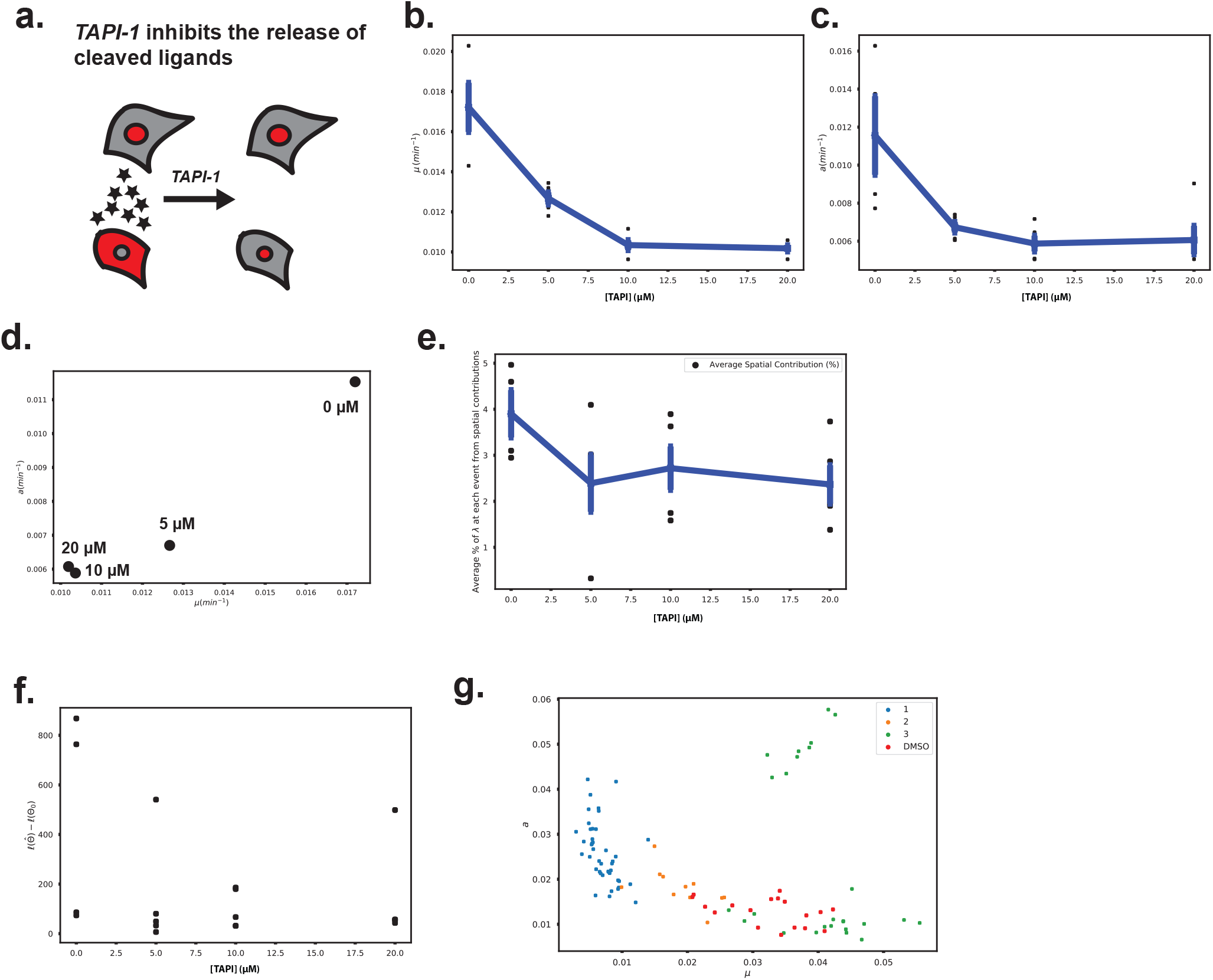
(a) Schematic of the effect of TAPI-1 on inter-cellular signaling. (b, c) Estimated parameters *μ* and *a* for keratinocytes as a function of TAPI-1 concentration. Black points represent individual wells. (d) Scatter plot of average estimates of *μ* versus *a* for each TAPI-1 concentration. (e) Average contribution of spatial signaling to pulses in each TAPI-1 concentration. (f) Difference in unnormalized log-likelihood of spatial model versus a spatial-free control model for all TAPI-1 concentrations and replicates. (g) Class 1, 2, and 3 drugs (6) on a plot of *μ* versus *a*.

**Table 3.**
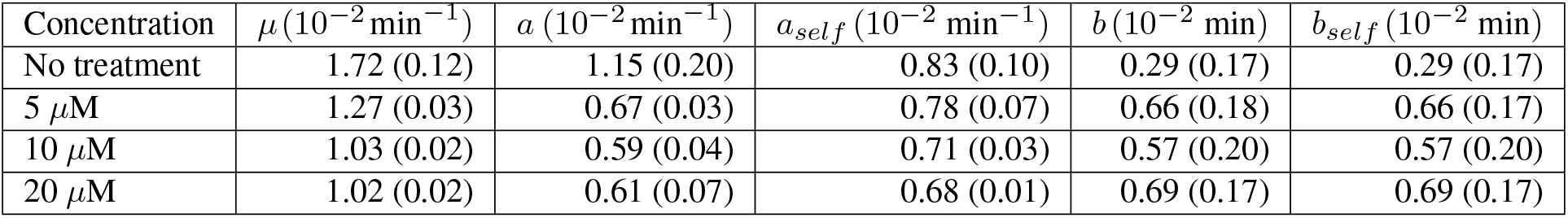
Mean estimated parameters and standard error of the mean (in parenthesis) for each TAPI-1 dose concentration.

**Fig. 4.**
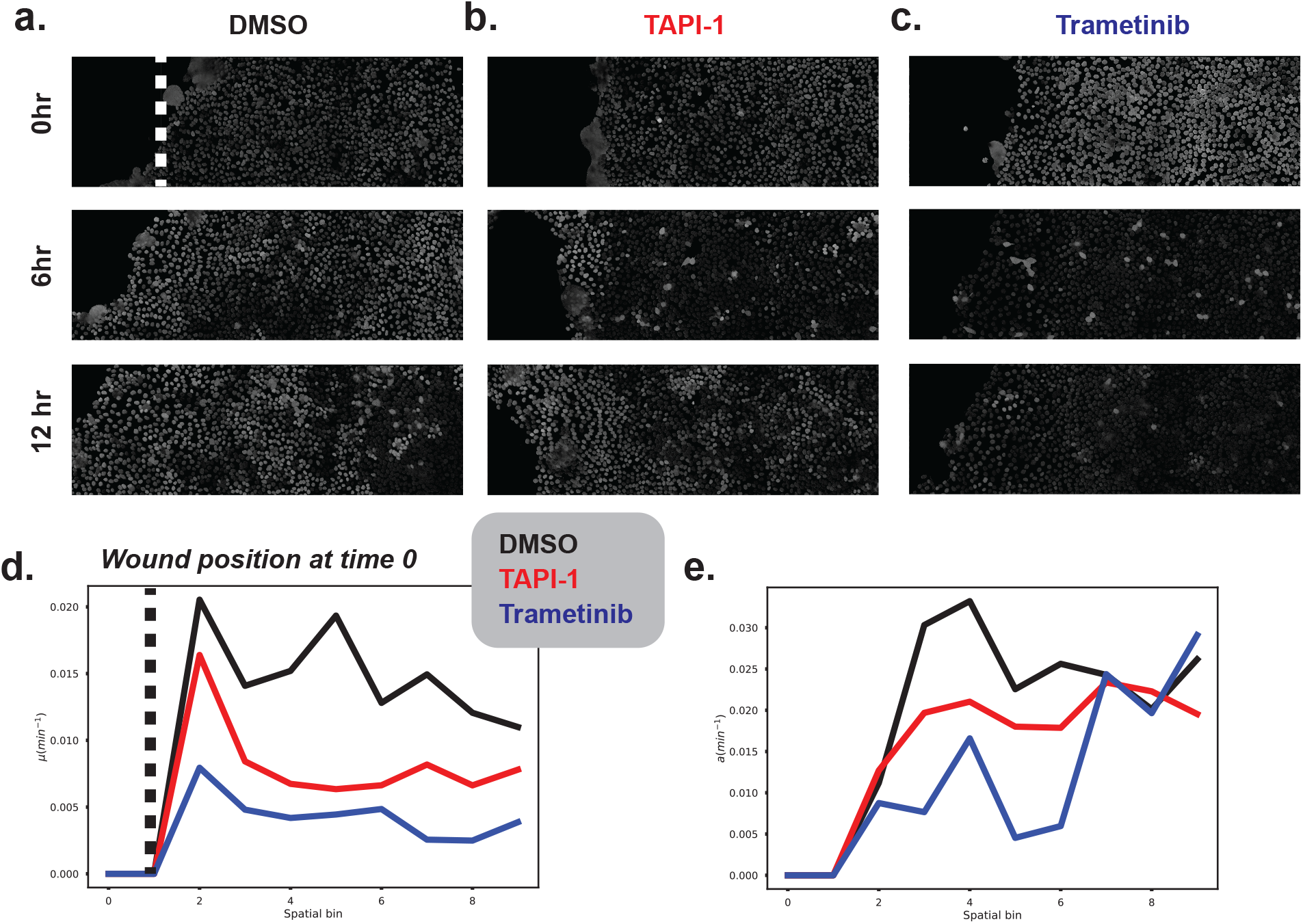
(a-c) Time course data of MDCK cells moving to heal a inflicted wound at 0, 6, and 12 hours post wounding. Dark cells denote low Erk activity, while light cells denote high Erk activity. (d) x-binning captures cells at different distances away from the site of a wound. *μ* estimated for bins at different distances away from the wound. (e) *a* estimates for spatial bins at different distances away from the wound. For both (d) and (e), black lines represent DMSO cells, the red lines represent TAPI-1 treated cells, and the blue lines represent trametinib-treated cells.

Linear regression between TAPI-1 concentration and the spatial parameter estimates 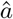, representing the presence of signaling at those TAPI-1 concentrations, shows a negative correlation (*β* = −0.0002, *p* ≤ 0.016), matching our prior biological knowledge. We also demonstrate the decrease in signaling associated with increase TAPI-1 dose using the likelihoods of the model. We compare the likelihood of the estimated model to a model where all pulses are due to autonomous signaling parameter *μ*, a.k.a *a* = 0. The estimated 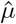 is the ratio of 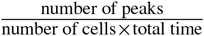. When we take the difference of the log-likelihoods, we observe that the difference between CPP and a control model decreases as TAPI-1 concentration increases (Figure 3f). This means that the CPP model is better at explaining the data relative to a fully autonomous model at low TAPI-1 concentrations. Alterna tively, we can calculate the contribution of the spatial, self-exciting, and autonomous influences at each peak. We observe that the average percentage of likelihood contribution from spatial signaling across peaks decreases as TAPI-1 concentration increases (Figure 3e). The ability of the CPP model to quantify this biological inhibition demonstrates its ability to analyze spatial signaling in complex systems.

We extend this analysis to evaluate the effects of a variety of drugs on spatial signaling in keratinocytes. Recent work (6) sought to quantify the effects of over 400 receptor tyrosine kinase inhibitors (RTKi) on endogenous keratinocyte Ras/Erk dynamics. We fit our CPP model on time series from treated cells from this study, with experiments across 432 drugs and 18 control DMSO samples, to determine whether the autonomous or spatial components, or both, are substantially affected by targeted RTK inhibition.

The original analysis of these data (6) divided drugs into three categories, which also took into account the “set point” or baseline level of Erk activity. Class 1 drugs reduced Erk activity to extremely low levels, corresponding to *μ* close to 0. Class 2 drugs, on the other hand, increased Erk activity to a high constant level. Since a pulsing event is defined as a local maxima in the Erk activity trace, these cells demonstrated very few pulsing events since the pathway could likely not be activated beyond this extremely high level. Since our model does not capture mean Erk activity, but rather the events where activity changes, we expect these drugs to have lower autonomous pulsing parameter *μ* in the CPP model. Finally, class 3 drugs increased Erk activity over time by increasing the pulse frequency (6), which would correspond to a higher estimated *μ* value in our model.

The CPP model finds differences across these three classes and in comparison to untreated DMSO controls (Figure 3g). We find that class 1 (*p* ≤ 8.2 × 10^−30^) and 2 (*p* ≤ 5.7 × 10^−6^) drugs have lower mean autonomous activity 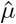 than DMSO controls. Class 3 drugs have higher autonomous activity 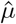 (*p* ≤ 0.0006). While class 1 and 2 drugs have spatial signaling parameter 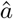 in a close range (approximately 0.01 − 0.04min^−1^), we find that class 3 drugs diverge between low signaling, 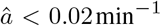 and high signaling behavior 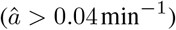. An interesting example is Pazopanib 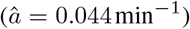, a class 3 drug that increases signaling relative to DMSO (mean = 0.013 min^−1^) that is known to target the membrane proteins such as RIPK1 and VEGFR (6, 40). Interactions with membrane proteins would be expected to be modulate inter-cellular signaling.

While estimates for autonomous parameter 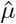 and signaling strength parameter 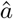 both decrease as TAPI-1 dose concentration increased, in the drug screen increasing 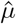 mostly corresponds to decreasing 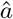. The divergence in class 3 drugs, however, indicates that both parameters are necessary to fully characterize the signaling system. One would expect that the addition of TAPI-1 would decrease the pulsing of the high 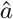 class 3 drugs but would leave the pulsing in low 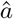 class 3 drugs mostly unaltered. The estimation of these distinct signaling parameters add nuance to our understanding of the cell response to various pharmacological agents beyond basic statistics such as frequency and duration of pulses.

### CPP quantifies trends in cell signaling in a wound healing context

Madin-Darby Canine Kidney (MDCK) cells move to close an artificially inflicted wound *in vitro* while expressing a live cell reporter of Ras/Erk activity (34). We analyzed these spatiotemporal data using TrackMate to obtain pulse information and cell positions from the cell images over time. We then fit our CPP model with these data. Cells were binned into their relative distance from the wound edge; the total width of the cell sheet from the inner edge of the field-of-view to the wound was split into ten bins, and the CPP model was fit for each bin (Figure 4a-e).

The estimated value of autonomous pulsing parameter 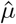 peaked next to the site of the wound in control (DMSO treated) cells (Figure 4d). On the other hand, the addition of TAPI-1 pre-wounding resulted in a much lower autonomous pulsing rate in this region (red curve, Figure 4d). Trametinib, an Erk inhibitor, decreased autonomous pulsing even further, as would be expected. The estimated value of inter-cellular signaling strength parameter 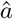 also decreased from DMSO to TAPI-1, and decreased further still in trametinib-treated cells (Figure 4e). Taken together, these results suggest, in line with previous work (34) that both cell-autonomous and cell-to-cell signaling effects occur with specific spatial organization in response to a wound, and can be abrogated to different extents through pharmacological inhibition. This suggests that both cell-autonomous and cell-to-cell Ras/Erk signaling may be important factors in allowing MDCK cells to close a wound. This may be why TAPI-1 and trametinib-treated cells fail to fully heal a wound over the full 12 hour time course (Figure 4b,c), where the control wound is fully healed at the 12 hour time point.

### CPP quantifies signaling-to-gene relationships using multi-channel learning

Engineered mouse keratinocytes that express a destabilized green fluorescent protein (dGFP) with a half-life of ~ 1 hour, under the control of the minimal promoter of the IEG cFos, were imaged for 24 hours. CPP allowed us to quantify the signaling between channels, Erk and Fos, with no prior information that Erk affects transcription of Fos. To measure this inter-channel signaling, we estimated the inter-cellular signaling parameters *a_ktr→gfp_* and *a_gfp→ktr_* using the multi-variate CPP model, and took the ratio of the two values for each field of cells imaged (Figure 5d). We were able to recapitulate the strong directed signaling of Erk to Fos transcription, as seen by the ratios of 3 and 2 for cells in growth and starved media, respectively (Figure 5e). On the other hand, testing several drugs from a previously published keratinocyte drug screen (6) showed a range of signaling behaviors (Figure 5e). Erk inhibition (UO126, Lapatinib) showed similar signaling in the presence of Erk-activating drugs GDC-0879 and SB590885, perhaps because the signaling between pulses of Erk signaling and pulses of GFP accumulation decreases when both are constantly on or constantly inhibited. On the other hand, tivozanib and pazopanib, which increase pulsing frequency *μ* (6), maintain a similar level of signaling and gene expression as cells grown in standard or starved conditions (Figure 5e).

**Fig. 5.**
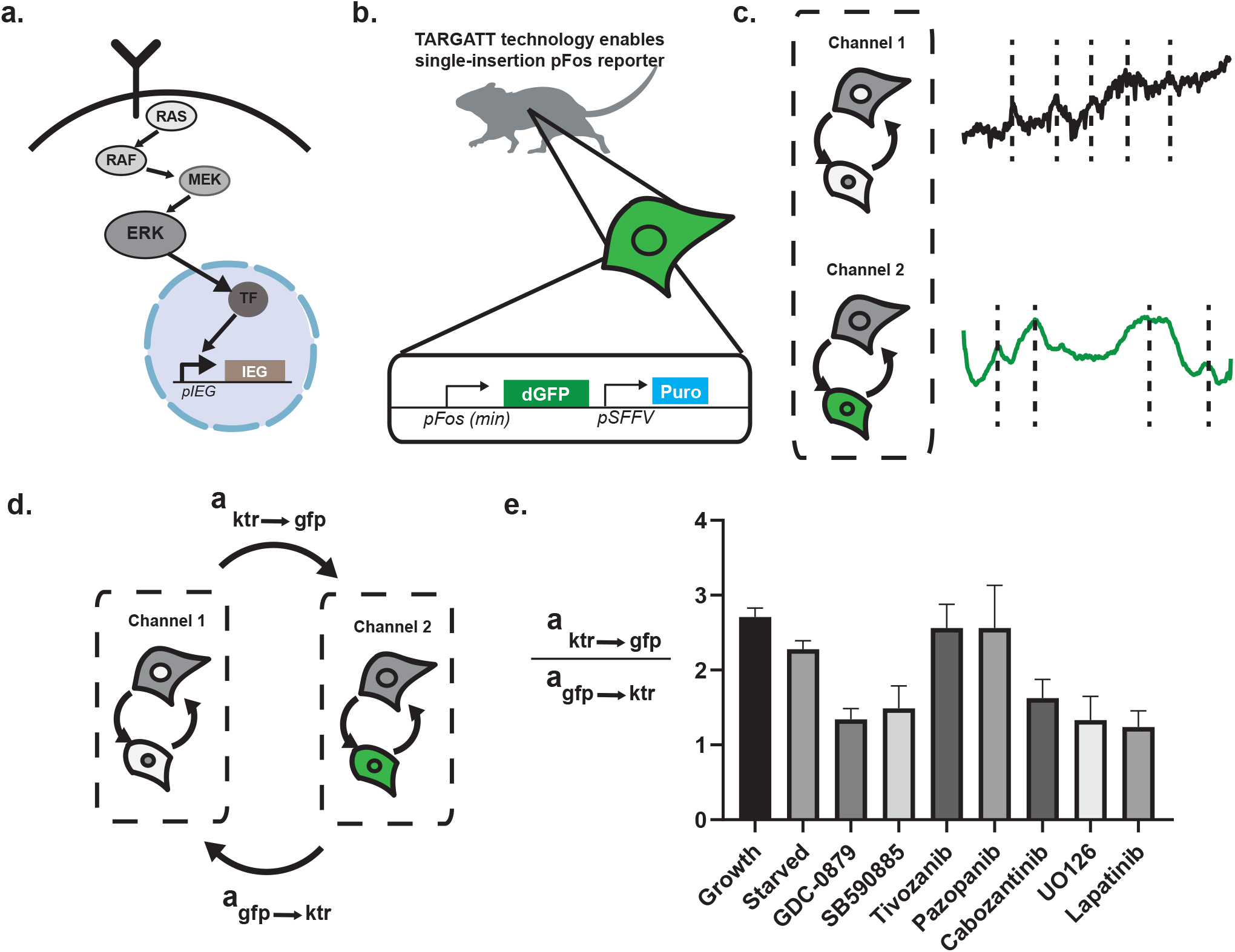
(a) The Ras/Erk pathway results in the transcription of canonical Immediate Early Genes. (b) TARGATT technology enables insertion of a Fos minimal promoter (pFos) into a single site in the genome, allowing for quantitative monitoring of pFos activation. (c) KTR and dGFP readout can be measured in the same cell. (d) CPP quantifies cross-channel directed signaling through *a_ktr→gfp_* and *a_gfp→ktr_*. (e) Pharmacological inputs modulate the strength of the association between Ras/Erk signaling and Fos gene expression dynamics.

## Conclusions

In this paper, we present a spatiotemporal model, the cellular point process (CPP), to capture pulsatile cell signaling events based on self-exciting Hawkes processes. Applying this model to processed cell imaging data across time, we estimate model parameters that quantify the strength of spatial and autonomous signaling in a multi-cellular system, even in the context of heterogeneous cell types, multiple signaling channels, or environmental conditions. We used these parameters to quantitatively compare systems, e.g., pre- and post-exposure to pathway-targeting drugs. We validated our model on simulated data and demonstrated its ability to capture known inhibition effects of TAPI-1 on spatial Erk coupling in keratinocytes. We then used the CPP model to interrogate the effects of different drug treatments on keratinocytes and were able to replicate the known effects on expression of three classes of drugs and extend knowledge of the effects of the drugs to cell signaling. We find that the estimation inter-cellular signaling parameters and autonomous pulsing parameters adds to our understanding of keratinocyte drug response. Lastly, we use the CPP model to capture heterogeneous cell signaling behaviors across distance from the wound frontier in response to wound healing.

The proposed CPP model leaves room for further development. Signaling mechanisms may have refractory periods, time after which the likelihood of an event is depressed, leading to a different kernel that can take on negative values. The kernel and the relationship between distance and signaling strength might be better modeled non-parametrically to account for differences in cell size, shape, and imaging scales. Currently, the model assumes that interactions are local and symmetric. However, systems such as wound healing may demonstrate global and directional behavior. The model of spatial interaction could be modified to vary over the region. Alternatively, the intensity of interaction could be a function of the vector distance between positions rather than a scalar distance. For longer histories of observations, we would also be interested in allowing these parameters to vary over time, possibly to detect possible switch-points between local and global signaling regimes.

The probabilistic aspects of the model could also be expanded to make the model fully Bayesian. Priors may be placed on the parameters to learn a maximum *a posterior* estimate or the posterior distribution over parameters. Hierarchical models may be developed to estimate smoothly varying model parameters across the space instead of binning cells by distance to the wound. Hierarchy could be used to learn shared parameters from multiple observations of the same condition such as repeats of the DMSO controls. Regardless, self-exciting point processes, and the CPP specifically, represent a powerful class of stochastic models that can be used to accurately and robustly quantify the spatial components of multi-cellular dynamics.

## ACKNOWLEDGEMENTS

We would like to thank Alexander Goglia for kindly providing data and annotations from his experimental drug screen (6).

